# Developmental variation in ovaries and kernels in a multi-kernelled rice variety- Jugal

**DOI:** 10.1101/2020.09.21.305995

**Authors:** Pallabi Saha, H.A. Murali, Bidhan Roy

**Author notes:** All authors contributed equally to this work.

## Abstract

The rice fruit or spikelet contains only one kernel. A handful traditional cultivars are available with more than one ovaries/kernels per spikelet. *Jugal*, a traditional rice cultivar from West Bengal, India possess multiple ovaries before flowering and multiples kernels per spikelet. In this endeavour, the single grained-spikelet varied from 0.00 to 46.26% per panicle. Single pistillated-spikelet was first reported on 8^th^ day before flowering and it was 2.63%. The number of spikelet bearing single ovary/kernel gradually increased till the panicle emergence and then till physiological maturity of grains. Maximum single kernelled-spikelet was recorded on 30^th^ day after panicle emergence (46.26%). The double ovary/kernelled-spikelet per panicle varied from 53.74% to 95.39%. High percentage of double pistillated-spikelet per panicle was recorded before heading and it was more than 90% or very near to 90%. Gradually the double pistillated-spikelet per panicle decreased with the advancement of developmental course of the panicle and continued to decrease till grain maturity. Lowest percentage (53.74%) of double kernelled-spikelet per panicle was observed on 30^th^ day after panicle emergence and it almost remained static till harvest of the crop. Triple pistillated-spikelet was 4.33% per on 10^th^ day before panicle emergence. On 9^th^ day before panicle emergence it was 2.33% and on 8^th^ day before panicle it was 1.00% only and subsequently, no triple pistillated-spikelet was observed till grain maturity. However, randomly one/two triple kernelled-spikelet was also reported. It would be useful if all the spikelets were doubled kernelled. The shape of kernels obtained from doubled kernelled-spikelets were slender, which has high demand among the urban and sub-urban consumers.

## Introduction

Rice is known to be the starchy cereal responsible for feeding two-thirds of the world population. Being the staple food across Asia, Africa and it is becoming evidently important in Africa and Latin America. Domestication and cultivation of rice is considered and ranked as one of the most important historic developments. The cultivation rate of rice is much higher than any other crop and the socio-economic, geographic conditions are quite vast. Through the course of cultivar selection since ancient time, a huge change in genetic and morphological architecture had took place towards desirable direction leading to creation of enormous genotypic variability. India is one of the richest hubs of rice diversity. The traditional cultivars were recorded, collected and studied for their desirable traits. The local landraces not only maintain genetic diversity but also can exceed many modern varieties in special characters and for cultivation in difficult soils.

Apart from the agronomically distinguishable characters which are preferred by the consumers and the farmers, some cultivars exhibit special traits which can be useful for further breeding or cultivation purposes or characterization for its distinctive features. The special traits exhibited are multiple spikelet developed from a single pedicel (e.g. Khejurchari), multiple carpels per ovary, multiple stigmas per gynoecium, and multiple kernels (2-3 mostly) per spikelet [1,2].

Rice is usually considered to be a crop yielding single grain, *i*.*e*., one spikelet will bear one kernel. However, the traditional variety of rice from West Bengal-the *Jugal* reveals that the cultivar bears multi-kernelled spikelets and it bears 2-3 kernels per spikelet. Taking into account the significance of multi-kernelled rice cultivar, the developmental variations of spikelet of *Jugal* was studied in this communication.

## Materials and methods

At panicle initiation stage, 700 main tillers (one main tiller from each hill) were tagged with blue coloured plastic tags bearing the date of tagging. Those tagged plants were used for study of developmental variations.

### Number of ovary/kernel per spikelet and stigma per ovary

Every day starting from panicle initiation, five panicles from the tagged plant were collected to study the number of ovary till flowering and then the number of kernel till harvestable maturity. Spikelet from collected panicles were split open with needle and forceps to study the number of ovaries. The split spikelets were observed under stereo-microscope and data were recorded on number of spikelets bearing multiple ovaries and single ovary. After fertilization, the ovary/kernel size were comparatively bigger and the observation on number of kernel per spikelet and number of stigma per ovary was taken in necked eyes.

### Weight of ovary/kernel

Average weight of ovary was studied on alternate days and started from 7^th^ day before panicle emergence. The same spikelets which were used for study of number of ovaries per spikelet were also used for recording observation on weight of ovary/kernel. As the size of the ovaries in the initial stage was very small, 100-ovaries were used for recording weight and after flowering 50-kernels were used for recording weight.

### Quantitative and qualitative characters

Observation of various qualitative and quantitative characters of *Jugal* were recorded at different stages of plant growth with appropriate procedures as outlined in the “Guidelines for the Conduct of Test for Distinctiveness, Uniformity and Stability on Rice (*Oryza sativa* L.)” published by Protection of Plant Varieties and Farmers’ Rights Authority (2007) [3], New Delhi, Government of India.

## Results

### Developmental course of ovaries/kernels

The single grained-spikelet varied from 0.00 to 46.26% per panicle (Fig 1). It was thought-provoking that there was no single pistillated-spikelet per panicle from panicle initiation to 9^th^ day before panicle emergence. Single pistillated-spikelet was first reported on 8^th^ day before flowering and it was 2.63% (Fig 2A). The number of spikelet bearing single ovary/kernel gradually increased till the panicle emergence and then till maturity of the grain. Maximum single kernelled-spikelet was on 30^th^ day after panicle emergence (46.26%). As the method of taking observation was based on destructive-sampling, negligible increase and decrease in percentage of single kernelled-spikelet were observed during the course of maturity of the grains.

**Fig 1.**
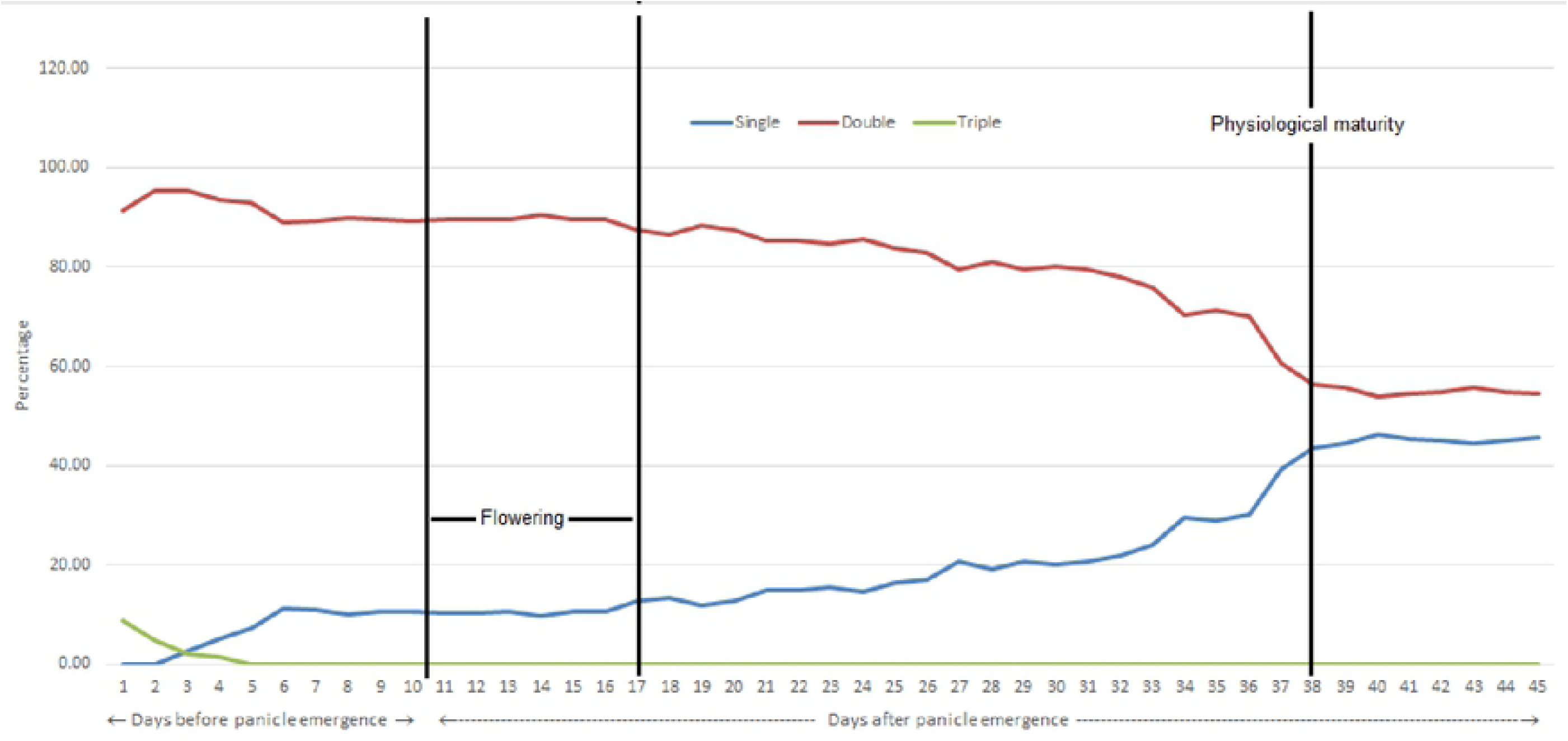
Developmental variations in kernel per spikelet of *Jugal* during the course of development of panicle and grain maturity.

**Fig 2.**
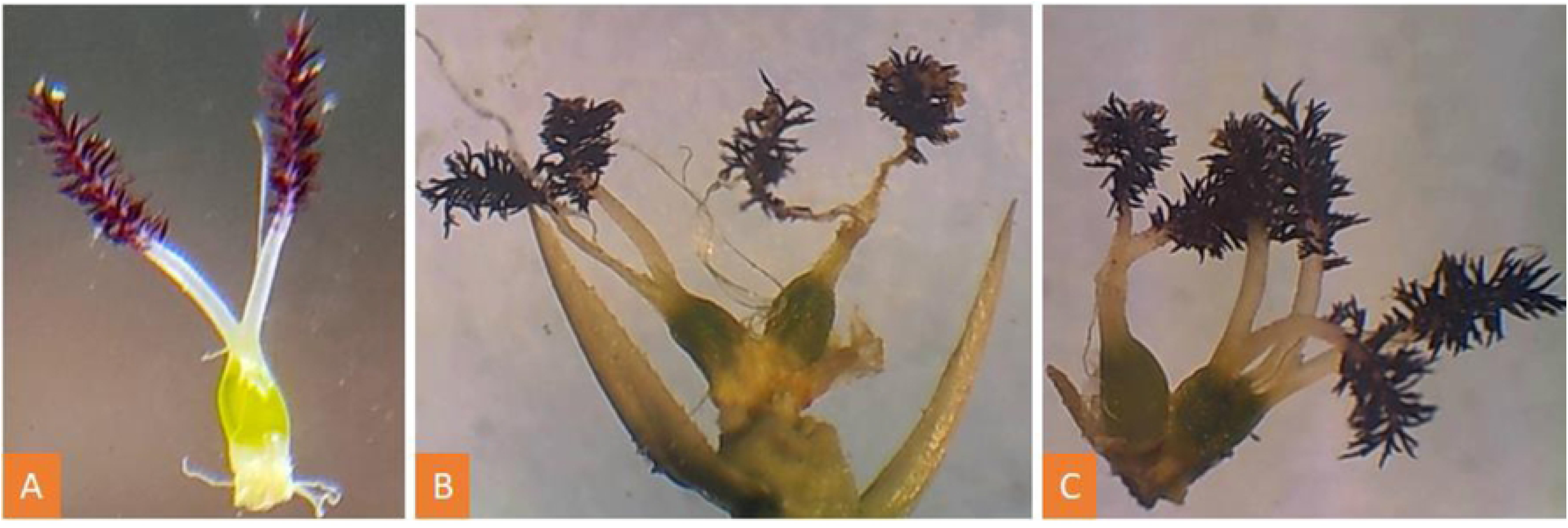
Spikelets with multiple-ovaries. **(A)** Spikelet with one ovary. **(B)** Spikelet with two ovaries **(C)** Spikelet with three ovaries.

The double ovary/kernelled-spikelet per panicle varied from 53.74% to 95.39% (Fig 1). High percentage of double pistillated-spikelet per panicle was recorded before emergence or heading and it was more than 90% or very near to 90% (Fig 2B). Gradually the double pistillated-spikelet per panicle decreased with the developmental course of panicle till grain maturity. Lowest percentage (53.74%) of double pistillated-spikelet per panicle was observed on 30^th^ day after panicle emergence and it almost remained static till harvest of the crop (Fig 1).

Triple pistillated-spikelet was observed only during initial stage of panicle formation. In this endeavour, 4.33% of total spikelets per panicle was observed on 10^th^ day before panicle emergence. On 9^th^ day it was 2.33% and on 8^th^ day it was 1.00% only and subsequently, no triple pistillated-spikelet was observed till grain maturity (Fig 1). However, randomly one/two triple kernelled-spikelet was reported which was not recorded in the Fig 1.

### Distribution of multi-kernelled spikelets in panicle

The distribution of multi-kernelled spikelets at harvestable maturity was not uniform in the entire panicle. About 70.60% of the multi-kernelled spikelets were found in the upper portion of the panicle, 23.80% were found in the middle portion of the panicle and 5.60% were found in the lower part of the panicle (Table 1). The causes of the low percentage of multi-kernelled spikelets in the lower portion of the panicle at harvestable maturity may be the malnutrition of spikelets at lower portion of panicle. In general, lower part of panicle showed higher number of chaffy grains as compared to upper portion.

**Table 1.** Distribution of double kernelled spikelets of *Jugal* in a panicle.

| Part of the panicle | Distribution of double kernelled spikelets in a panicle |
| --- | --- |
| Upper portion | 70.60% |
| Middle portion | 23.80% |
| Lower portion | 5.60% |

### Weight of ovary/kernel

Gradually the weight increased during the course of panicle development and grain filling (Fig 3). The pictorial representation of double ovaries and kernels were given in Fig 4. Ovary weight of single ovary-spikelet on first day of panicle emergence was 0.0102 g, whereas single ovary weight of double ovaries-spikelet on first day of panicle emergence was 0.0056 g. In most of the cases, the average weight of ovary of single ovary/kernelled-spikelets was comparatively lower than the added weight of two ovaries/kernels of corresponding double ovaries/kernelled-spikelets (Fig 3). The kernel weight increased rapidly in between the panicle emergence and till completion of physiological maturity of grain. Test weight of single kernelled-spikelet at harvestable maturity was 29.90 g (Table 2), however the test weight at physiological maturity was 33.50 g (Fig 3). The decrease in test weight at harvestable maturity was due to reduction in kernel moisture content at harvestable maturity.

**Table 2.** Quantitative and qualitative characters of *Jugal*.

| Sl. No. | Characters | Status |
| --- | --- | --- |
| 1. | Coleoptile colour | White |
| 2. | Basal leaf sheath colour | White |
| 3. | Intensity of green colour of leaf | Medium |
| 4. | Anthocynin colouration of leaf | Absent |
| 5. | Leaf sheath anthocynin colouration | Absent |
| 6. | Pubescence of blade surface of leaf | Very Strong |
| 7. | Auricles | Present |
| 8. | Anthocynin colouration of auricles | Colourless |
| 9. | Leaf collar | Present |
| 10. | Anthocynin colouration of leaf collar | Absent |
| 11. | Ligule | Present |
| 12. | Shape of ligule | Split |
| 13. | Colour of ligule | White |
| 14. | Length of leaf blade | Long |
| 15. | Width of leaf blade | Medium |
| 16. | Culm attitude | Erect |
| 17. | Time of heading (50% plants with panicles) | Late (122 days) |
| 18. | Attitude of flag leaf blade (early observation) | Erect |
| 19. | Density of pubescence of lemma | Medium |
| 20. | Male sterility | Absent |
| 21. | Anthocyanin colouration of keel of lemma | Absent |
| 22. | Anthocyanin colouration of area below apex of lemma | Absent |
| 23. | Anthocynin colouration of apex of lemma | Absent |
| 24. | Colour of stigma | Dark Purple |
| 25. | Stem thickness | Medium (0.46 cm) |
| 26. | Stem length (excluding panicle; excluding floating rice) | Very Long (160 cm) |
| 27. | Anthocyanin clouration of nodes | Absent |
| 28. | Anthocynin colouration of internodes | Absent |
| 29. | Length of main axis of panicle (cm) | Medium (24.70 cm) |
| 30. | Attitude of blade of flag leaf (late observation) | Semi - erect |
| 31. | Panicle curvature of main axis | Deflexed |
| 32. | Number of panicles/plant | Medium |
| 33. | Colour of tip of lemma | Purple |
| 34. | Colour lemma & palea | Straw |
| 35. | Awns | Present |
| 36. | Colour of awns (late observation) | Straw |
| 37. | Length of longest awn | Medium |
| 38. | Distribution of awns | Upper half only |
| 39. | Presence of secondary branching in panicle | Present |
| 40. | Secondary branching in panicle | Strong |
| 41. | Attitude of branches in panicle | Erect to semi erect |
| 42. | Panicle exertion | Well exerted |
| 43. | Time of maturity (days) | Very Late (>160 days) |
| 44. | Leaf senescence | Medium |
| 45. | Colour of sterile lemma | Straw |
| 46. | Weight of 1000 fully developed grains (g) | High (30 g) |
| 47. | Grain length (mm) | Short (7.35 mm) |
| 48. | Grain width (mm) | Medium (3 mm) |
| 49. | Decorticated grain length (mm)- Single kernelled spikelet | 5.73 mm |
| 50. | Decorticated grain length (mm)- Double kernelled spikelet | 5.65 mm |
| 51. | Decorticated grain width (mm)- Single kernelled spikelet | 2.81 mm |
| 52. | Decorticated grain width (mm)- Double kernelled spikelet | 1.62 mm |
| 53. | Decorticated grain shape – Single kernelled spikelet | Short Bold |
| 54. | Decorticated grain shape – Double kernelled spikelet | Medium Slender |
| 55. | Decorticated grain colour | Dark Brown |
| 56. | Decorticated grain aroma | Absent |

**Fig 3.**
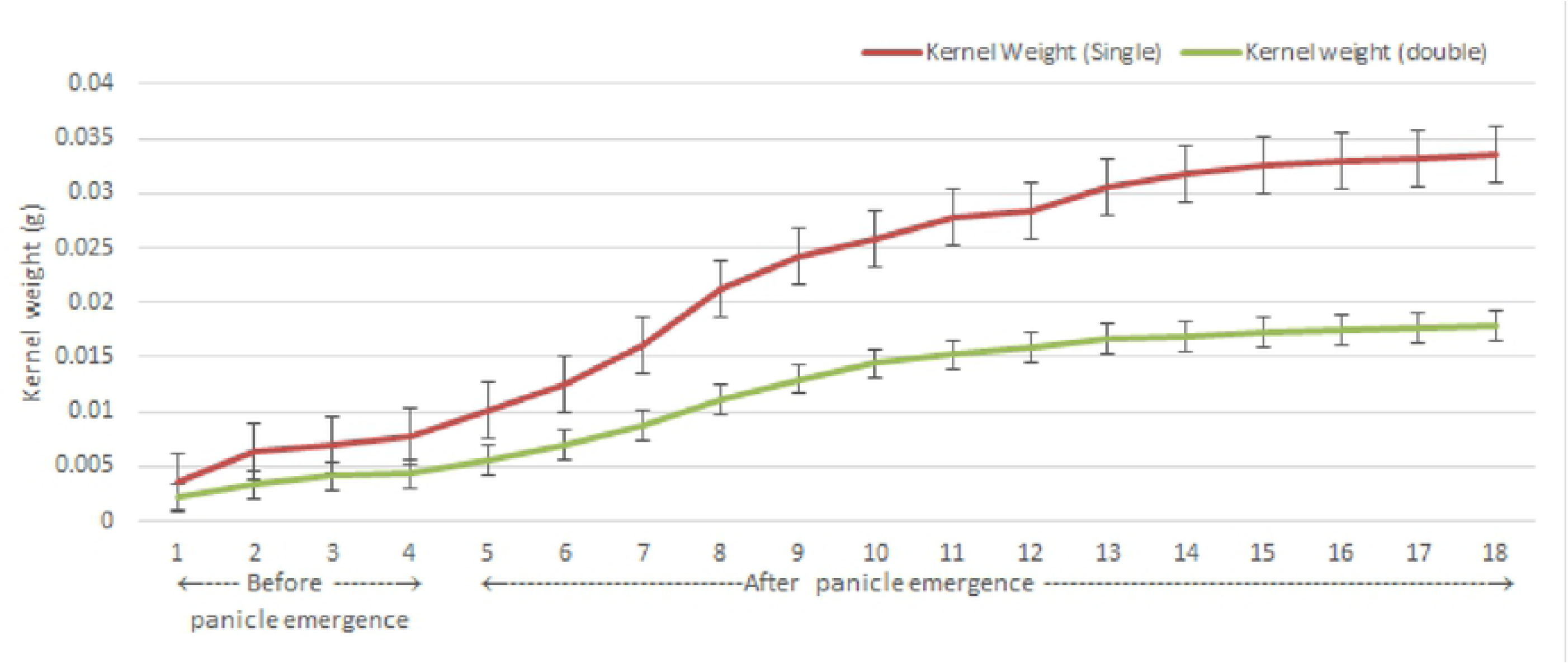
Weight of ovary/kernel in developmental course of ovary formation and kernel growth.

**Fig 4.**
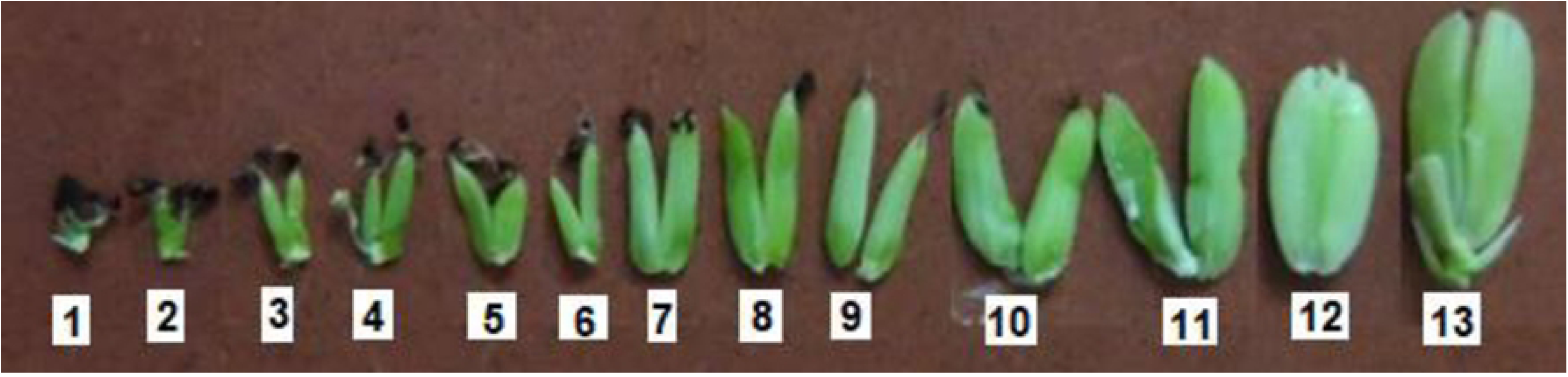
Developmental variations in shape and size of double kernelled spikelets of *Jugal*. 1-7, Before fertilization. 8-13, after fertilization.

### Number of stigma per ovary

In a normal rice floret, there are two bifid plumose (feathery) stigmas in the gynoecium system with short and thick style (Fig 5A). Rice ovary is superior with one ovule. The number varied from 2 (normal gynoecium, Fig 5A) to 5 (Fig 5D) stigmas per ovary. Multiple stigma per ovary was reported in this study (Fig 4.9). A spikelet was found with three ovaries and total of nine stigmas (Fig 5D).

**Fig 5.**
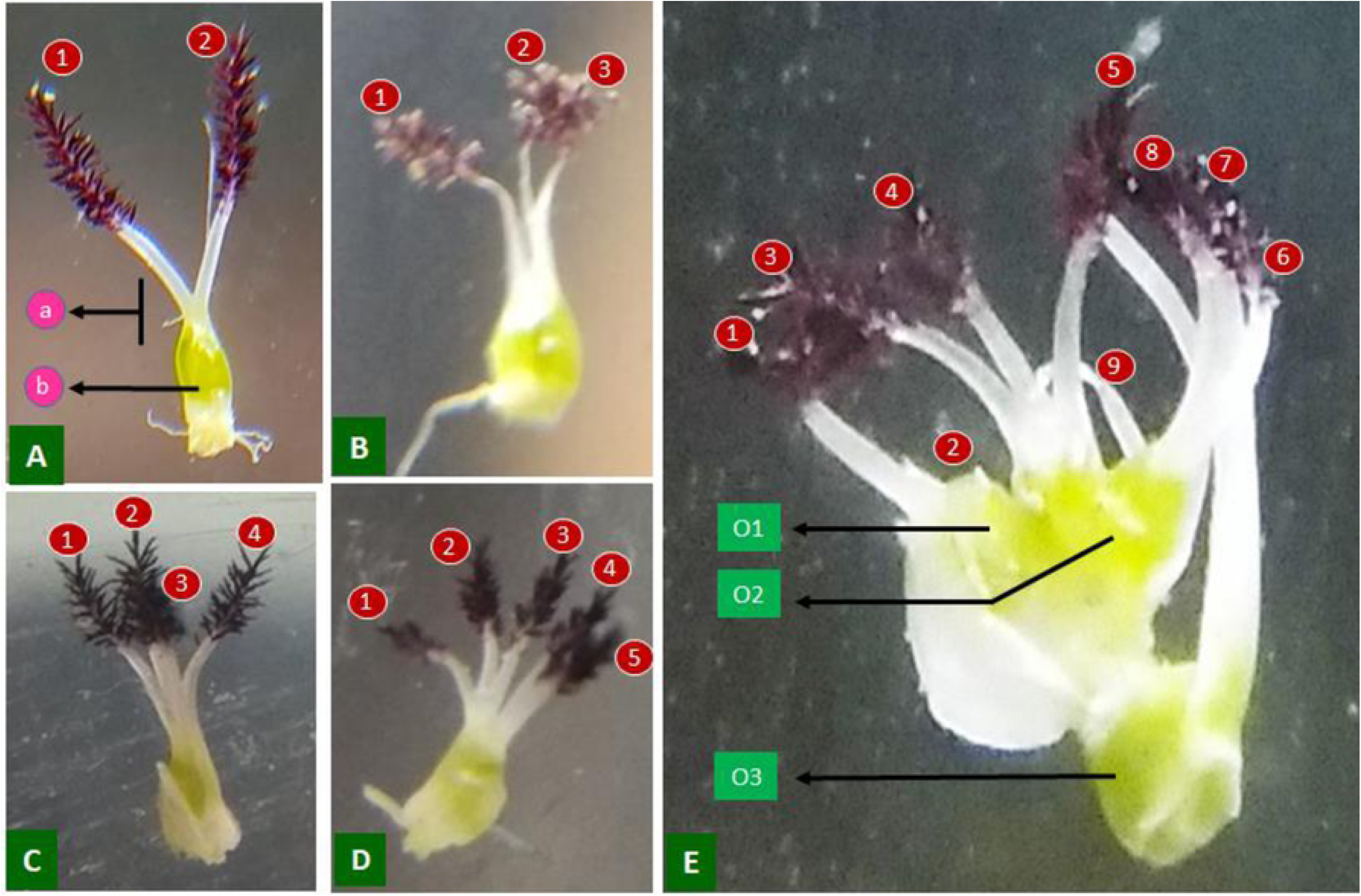
Variation in number of stigma in floret of Jugal. (**A**)Normal gynoecium of *Jugal* with two bifid stigma, short style and one ovary, ***(a)*** Short and thick style, ***(b)*** Mono-pistillate ovary. (**B)** Ovary with three stigma. (**C)** Ovary with four stigma. (**D)** Ovary with five stigma (**E)** Triple ovaried spikelet and total of nine stigma within the spikelet, ***(O1)*** Ovary with four stigma [1, 2, 3 & 4], ***(O2)*** Ovary with three stigma [5, 6 and 7], ***(O3)*** Ovary with two stigma [8 & 9].

### Quantitative and qualitative characters of *Jugal*

The quantitative and qualitative characters data were presented in Table 2. The cultivars was very tall (160.00 cm), long duration (163 days), medium tillering ability (10-15 tillers per plant), medium panicle length (24.70 cm), high test weight (30.00 g), low number of filled grains per panicle (157/panicle), adequate number of chaffy grains per panicle (43/panicle) and comparatively high sterility (27.37%), low grain yielder (2.50 t/ha).

Some of the unique characteristics of the cultivar were purple coleoptile colour, dark purple stigma (Fig 5), occasional presence of multiple stigma on single ovary (Fig 5), purple coloured long awn, frequently presence of multiple ovaries (Fig 2&4) and multiple kernels per spikelet (Fig 2&4), red aleurone layer. The cultivar was highly photoperiod sensitive, it can be grown only during *Kharif* season.

The grain characters were presented in Table 2. Straw coloured lemma and palea. Decorticated grain length and breadth from single kernelled spikelet was 5.73 mm and 2.81 mm, respectively. The decorticated grain length and breadth of kernel from double kernelled spikelet was 5.65 mm and 1.62 mm, respectively. The decorticated grains from single kernelled spikelet was classified as short bold and the decorticated grains from double kernelled spikelet was classified as medium slender. Aleurone layer of decorticated grain was red coloured. The undehusked grain length and breadth was 7.35 mm and 3.00 mm, respectively.

## Discussion

Rice ovary is superior, one celled with a single ovule. However, some of the previous authors had reported more than one ovules per ovary [4]. Multiple kernels also had been observed in few of the traditional varieties of rice [1, 5-7]. Rice gynoecium usually bears only one ovary with two stigma and a single carpel. Formation of the two ovaries in a spikelet is very unique trait, because one-floret per spikelet is the characteristics of the *Oryza* genus and is strictly regulated [8]. Multiple ovary within a single spikelet is a very uncommon and rarely noted character in rice floret morphology. Very few report are available on study of multiple ovary in rice. Most of the previous researchers studied the proportion of many kernelled spikelets in a panicle at grain maturity. In this venture, the occurrence of single, double and triple kernelled-spikelets per panicle at harvestable maturity was 46.26%, 53.74% and 0.00%, respectively. Tin and Kang [9] studied the flower structure of rice cultivars, namely, ‘Yu Sze 3’ and ‘Compound Rice’ and observed the variation in number of ovaries in a floret. Jeremy Cherfas [10] also reported multi-kernelled spikelet in ‘Laila Majnu’ (IRRI gene bank accession No. IRGC 59101) and ‘Amaghauj’ local cultivars of rice in Nepal. A group of farmers in Bangladesh were conserving a unique rice variety of rice, ‘Biram Sundori’. It has two (sometime three) kernels in a spikelet [11].

All those authors did not mentioned any proportion of many pistillated-floret per panicle. Roy *et al*. [6] also suited the presence of many kernelled spikelets in Jugal. Chakrabarty *et al*. [12] reported the occurrence of single, double and triple-kernelled spikelet were 53.7%, 41.3% and 5.0%, respectively in a panicle. Occurrence of single, double and triple kernels per spikelet in *Jugal* as had been observed by Roy and Surje [1] was 53.9, 42.2 and 3.9%, respectively. Gulam and Chanda [13] observed 50.28% single seeded, 49.30% double seeded and (0.42%) triple seeded spikelets per panicle. Contrary, our present findings did not match with findings of earlier researchers finding of occurrence proportion of single, double and triple kernelled-spikelets per panicle in *Jugal*. In our study, there was no triple kernelled-spikelet at harvestable maturity of grains. In the collection of Debal Deb, a cultivar ‘Sateen’ have three kernel per spikelet. This is a unique trait of three kernels per spikelet of Sateen (*O. sativa* var. *indica*) have been registered at National Bureau of Plant Genetic Resources (NBPGR) by Deb [14].

Abnormalities in number of stigma per stigma was also observed. The number varied from 2 (normal gynoecium) to 5 stigmas per ovary. Abnormal pistil with four stigmas in the *fon4-1* flower was observed by Chu *et al*. [15]. They also observed pistils had stigmas with 3-8 stigma branches. Increase in number of stigma may be the fusion of two or more carpels [15]. The spikelet meristem of *Ubi1:AtJMT* (jasmonic acid carboxyl methyltransferase gene) plants was enlarged, the number of spikelet organ primordia was altered, and the extra organ structures were modified in appearance [16]. They reported *Ubi1:AtJMT* plants with extra stigma branch. Zhang *et al*. [17] found aberrant pistils of *Epi-df* plants. As per their finding, about 8.5% of flowers of *Epi-df* plants had more than six stamens.

As per the findings of Suzaki *et al*. [18], mutations in *FON2* caused enlargement of the floral meristem, resulting in an increase in the number of floral organs keeping the vegetative and inflorescence meristems almost normal. Molecular cloning by Suzaki *et al*. [18] revealed that *FON2* encodes a small secreted protein, containing a *CLE* domain that is closely related to *CLAVATA3* in *Arabidopsis thaliana. FON2* transcripts are localized at the apical region in all meristems in the aerial parts of rice plants, showing an expression pattern similar to that of *Arabidopsis CLV3*.

In the variety ‘Plena’ of *Oryza sativa* Linn had recorded as many as seven ovaries by Datta and Paul [4]. They also reported more than one ovules per ovary in Plena. However, they found that after withering of stamen, only two ovaries continued to grow normally. Infrequently one of those failed to grow as fast as the other. As per our finding, this caused the increased in number of single grained spikelets per panicle during the progress of maturity of the grains. In some cases (about 50% of the spikelets in a panicle) both the ovaries grew equally leading formation of double kernelled rice spikelets. Inner faces of each of the kernel were flat. This description is agreement with the findings of Datta and Paul [4]. In case of three kernelled spikelets, always one kernel was much smaller than other two (Fig 6i). Middle kernel was flat at both sides, other two kernels were flat on the inner sides. Sometimes, one kernel remained very under developed and thin.

**Fig 6.**
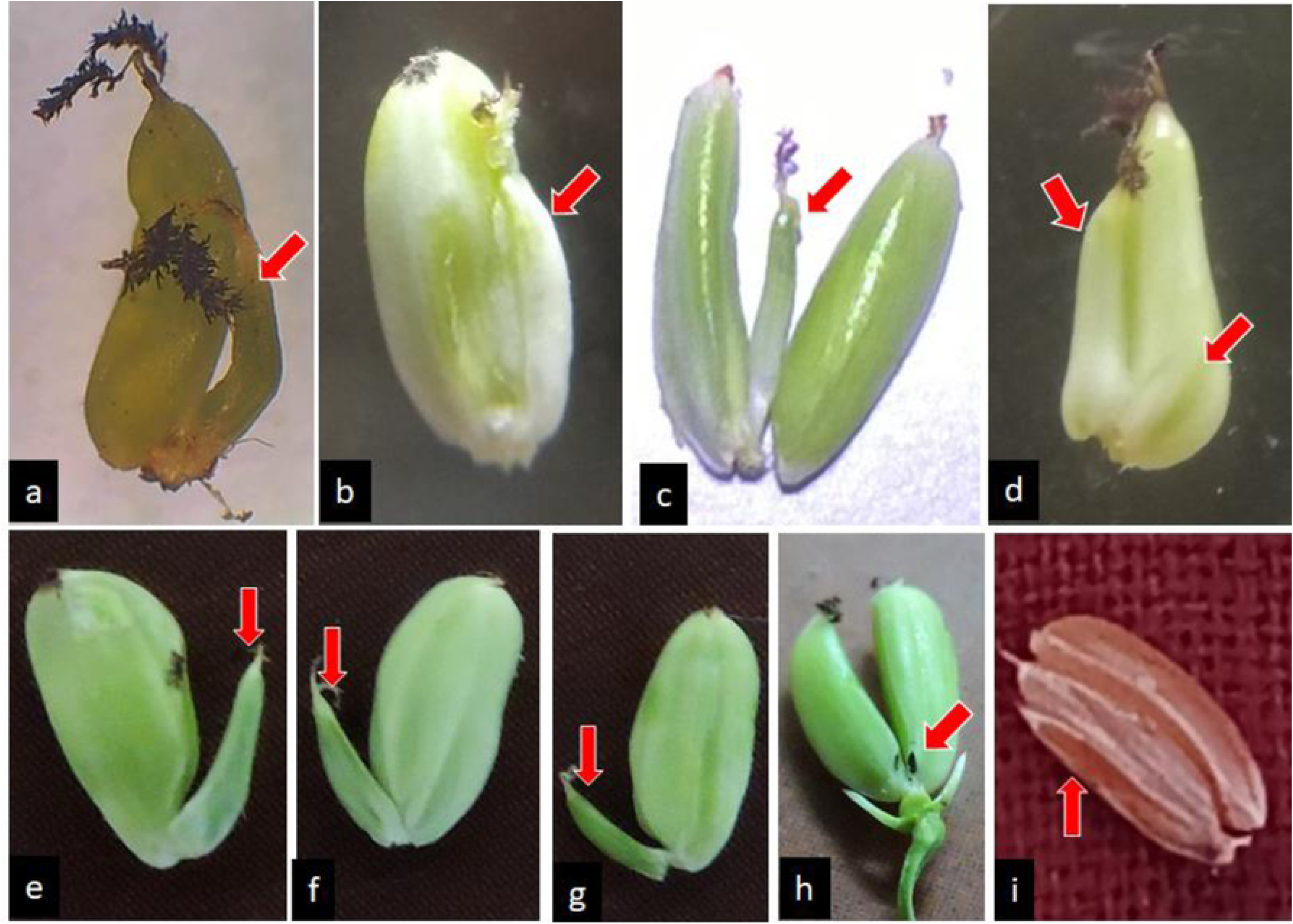
Degeneration of ovary or kernel during the course of ovary development and grain maturation. **Arrowhead** is the degenerating ovary/kernel. (**a, b, e, f & g**) Two pistillated spikelets. Ovary at one side (arrowhead) were underdeveloped as compared to the other ovaries. During the course of development, these underdeveloped ovaries will degenerate and the spikelets will turn to single kernelled spikelets. (**c & d**) Three pistillated spikelet. Arrowhead ovaries were underdeveloped as compared to the other two ovaries. During the course of development, these underdeveloped will degenerate and the spikelet will turn to a two kernelled spikelet. (**h**) Initially this spikelet was three pistillated spikelet. The scar at arrowhead indicated that one ovary already degenerated and it turned to double kernelled spikelet. **i**, Triple kernelled pikelet at harvestable maturity. Arrowhead kernel remained as rudimentary and it may be due to malnourishment or scarcity of space within the spikelet. (**a-d)** Degeneration of ovaries may be due to non-fertilization of those particular ovaries, because they are degenerating immediately after completion of flowering of panicle. (**e-g**) Degeneration of kernels may be due to malnourishment or by the pressure of fast growing other kernel(s) that created scarcity of space in the spikelet.

Old literature on developmental anatomy accomplishes that rice spikele was basically three-grained, two of which have become vestigial (“sterile lemma”). It would be interesting to note that the multi-grained spikelets are may be a reversion to primitive type of rice, or a new splitting of the central grain [11].

As per the conclusion of the findings of Priya *et al*. [19], “In normal cultivated rice and in *O. rufipogon* the meristematic activity in a spikelet stops after the production of the gynoecium, but in *Jugal*, the meristematic activity continues after fertilization, resulting in additional rudimentary pistil along with the mature normal one. But as the dimension is almost fixed for each spikelet, the mature kernels become reduced in size”. The findings of Priya *et al*. [19] is contradictory to our findings. We have observed multiple ovaries in a spikelets even well before fertilization, i.e. 10 days before panicle emergence. So, the hypothesis provided by Priya *et al*. [19] that “the meristematic activity continues after fertilization” may not work.

Most of the previous authors studied the proportion of many kernelled spikelets in a panicle at grain maturity. But, our study data on multi-pistil/kernelled-spikelets per panicle are available from the countable stage (10^th^ day before flowering) to harvestable grain maturity (Fig 1). Initially all the spikelets were with multiple kernelled. Maximum percentage of double pistillated-spikelet per panicle was recorded before heading of panicle and it was more than 90% or very near to 90%. Gradually the double pistil/kernelled-spikelet per panicle decreased with the developmental course of panicle till grain maturity. The probable cause of reduction of number of multi-pistil/kernelled-spikelets per panicle are being discussed below.

### Degeneration of ovary

During the course of ovary development and before anthesis degeneration of ovary has been reported (Fig 6a-h). The exact cause of degeneration of the ovary was not studied.

### Unfertilized ovary

The percentage of double ovary/kernelled-spikelets remained almost constant during flowering (from first flowering in a panicle to completion of flowering it takes about 8-10 days) and it decreased fast after completion of flowering. This decrease may be due to degeneration of unfertilized ovary (Fig 1) leading to increase the number of single kernelled spikelets in a panicle.

### Ill-filled kernelled or malnourished kernel

During the course of grain filling and nourishment, some of kernel may get malnourished (Fig 6). Those malnourished kernel gradually degenerate leading to increase the number of double to single kernelled spikelets (Fig 6a,b,e,f&g) or triple to double kernelled spikelets (Fig 6c,d&h) in a panicle.

### Limited space inside the spikelet

The shape of the kernel is decided by the shape of the lemma and palea. In case of multiple kernelled spikelets, under developed kernel or malnourished kernel may degenerates by the pressure of fast growing other kernel(s) that created scarcity of space in the spikelet as the space inside a spikelet is limited (Fig 6h) or almost fixed [19].

### Conclusion

This is the first time we are reporting the probable causes of reduction of multi-kernelled spikelets in a panicle in multi-kernelled rice cultivars in general and *Jugal* in particular. Already we mentioned that no previous researchers studied the number or percentage of multi-kernelled spikelets during the course of the panicle development and grain maturity. Previous authors recorded only the proportion of many-kernelled spikelets in a panicle at grain harvestable maturity. Research may be continued to find out the scientific was to retain the double kernel in all the spikelets in a panicle. Two kernels per ovary lead production of slender decorticated grains which has high consumers’ preference in urban and sub-urban areas.

## References

1. Roy B, Surje TD. Some special characteristics of farmers’ varieties of rice (*Oryza sativa* L.) for testing of distinctiveness. Indian J. Plant Genet. Resour. 2016; 29(2): 163–169.

2. Surje D, Kumar V, Roy B. Grouping of farmers’ varieties of rice (*Oryza sativa* L.) collected from West Bengal and adjoining states. Indian J. Plant Genet. Resour. 2018; 31(3): 251–259.

3. PPV & FRA. Guidelines for the conduct of test for DUS on rice (Oryza sativa L.). Protection of Plant Varieties and Famrer’s Right Authority (PPV & FRA), Government of India, New Delhi. 2007.

4. Datta RM, Paul AK. Morphology of the ovary of *Oryza Sativa* Linn. Var. *Plena* (“Double Rice”). Science. 1951; 113(2939): 491.

5. Chakravorty A, Ghosh PD. Characterization of Landraces of rice following DUS guidelines. Res Plant Biol. 2012a; 2(6): 30–40.

6. Roy B, Surje TD, Mahato S. Biodiversity of farmers’ varieties of rice (*Oryza sativa* L.) at repository of Uttar Banga Krishi Viswavidyalaya: A reservoir of important characters. The Ecoscan. 2013; 4: 145-151.

7. Das SP, Deb D, Dey N. Micromorphic and molecular studies of floral organs of a multiple seeded rice (*Oryza sativa* L.). Plant Mol. Biol. Rep. 2018; 36: 764–775.

8. Tanaka W, Toriba T, Yoshihiro OY, Hirano HY. Formation of two florets within a single spikelet in the rice *tongari-boushi1* mutant. Plant Signaling Behavior. 2012; 7(7): 793–795.

9. Tin PT, Kang CN. Morphological observations on many kerneled grains in rice. Acta Botanica Sinica. 1960; 9(1): 105.

10. Jeremy Cherfas. Triple-grained rice new. Available from: https://agro.biodiver.se/2012/02/triple-grained-rice-news/. 2012

11. Anonymous. Double Grained Rice in Bangladesh. Available from: http://agro.biodiver.se/2012/02/rice-morphological-diversity-1-bloggers-0/?utm_source=feedburner&utm_medium=email&utm_campaign=Feed%3A+AgriculturalBiodiversityWeblog+%28Agricultural+Biodiversity+Weblog%29). 2015.

12. Chakrabarty SK, Joshi MA, Singh Y, Maity A, Vashisht V, Dadlani M. Characterization and evaluation of variability in farmers’ varieties of rice from West Bengal. Indian J. Genet. Plant Breed. 2012b; 72(2): 136–142.

13. Gulam SAKM, Chanda SC. Morphology of a multiple seeded rice (Oryza sativa L.) cultivar of Bangladesh. Available from: https://www.researchgate.net/publication/328450936. 2018.

14. Deb D, Bhattacharya D. Two unique landraces from West Bengal. Seed Res. 2009; 37: 152–155.

15. Chu H, Qian Q, Liang W, Yin C, Tan H, Yao X, Yuan Z, Yang J, Huang H, Luo D, Ma H, Zhang D. The *FLORAL ORGAN NUMBER4* gene encoding a putative ortholog of Arabidopsis *CLAVATA3* regulates apical meristem size in rice. Plant Physiology. 2006; 142: 1039–1052.

16. Kim EH, Kim YS, Park SH, Koo YJ, Choi YD, Chung YY, Lee IJ, Kim JK. Methyl jasmonate reduces grain yield by mediating stress signals to alter spikelet development in rice. Plant Physiol. 2009; 149: 1751–1760.

17. Zhang L, Cheng Z, Qin R, Qiu Y, Wang JL, Cui X, Gu L, Zhang X, Guo X, Wang D, Jiang L, Wu C, Wang H, Cao X, Wan J. Identification and characterization of an Epi-Allele of *FIE1* reveals a regulatory linkage between two epigenetic marks in rice. The Plant Cell. 2012; 24: 4407–4421.

18. Suzaki T, Toriba T, Fujimoto M, Tsutsumi N, Kitano H, Hirano HY. Conservation and diversification of meristem maintenance mechanism in *Oryza sativa*: Function of the *FLORAL ORGAN NUMBER2* gene. Plant Cell Physiol. 2006; 47(12): 1591–1602.

19. Priya A, Das SP, Goswami S, Adak MK, Deb D, Dey N. An exploratory study on allelic diversity for five genetic loci associated with floral organ development in rice. American J. Plant Sci. 2015; 6: 1973–1980.

